# Association between neurofibromatosis type 1 and cerebrovascular diseases in children: a systematic review

**DOI:** 10.1101/2020.10.09.332841

**Authors:** Beatriz Barreto-Duarte, Fabiana H. Andrade-Gomes, María B. Arriaga, Mariana Araújo-Pereira, Juan Manuel Cubillos-Angulo, Bruno B. Andrade

**Author notes:** Corresponding authors (BBA), (BBD). B.B.D and F.H.A.G equally contributed to the work.

## Abstract

**Background:** Neurofibromatosis type 1 (NF-1) is an autosomal dominant disease that affects one in every 3000 individuals. This disease can present a wide range of clinical manifestations, ranging from skin abnormalities to severe vascular changes. Although little recognized, cerebrovascular diseases (CVD), often present since childhood and diagnosed late, may have clinical manifestations ranging from headache and cognitive deficits to aneurysm rupture causing death. Thus, the CVD play an important role in the clinical manifestations, the severity of the condition and the prognosis of patients with NF-1. This systematic review aims to summarize the body of evidence linking NF-1 and CVD the in children.

**Methods:** Two independent reviewers performed a systematic review on the PubMed and EMBASE search platforms, using the following key terms: “neurofibromatosis type 1”, “recklinghausen disease”, “children”, “adolescents”, “stroke”, “moyamoya disease”, “vascular diseases”, “cerebrovascular disorders”, “aneurysm” and “congenital abnormalities”. Studies focused on assessing the development of CVD in children with NF-1 were included.

**Results:** Seven studies met the inclusion criteria. Twelve different clinical manifestations have been associated with cerebrovascular changes in children with NF-1; 44,5% of diagnosed patients were asymptomatic.

**Conclusion:** The available evidence suggests that cerebrovascular diseases are related with the progression of NF-1, even in the absence of a clear clinical manifestation. In addition, better prognosis was observed when imaging tests were performed to screen for cerebrovascular changes. This generated early interventions and consequently more favorable outcomes.

## INTRODUCTION

Neurofibromatosis type 1 (NF-1) is an autosomal dominant, chronic and progressive disease [1] with an incidence of approximately 1:3000 individuals and almost half of the cases of hereditary origin (familial NF) [2]. The alteration in the *NF1* gene, located at chromosome 17 q11.2, is responsible for the inability to synthesize the neurofibromin cytoplasmic protein, which acts as a modulator in cell growth and differentiation since uterine life, and which is expressed in the nervous system, endothelium and smooth muscle cells of blood vessels. The gold-standard diagnosis can be achieved through molecular genetic testing, which is related to high cost and low availability in the public health system [3] Frequently, diagnosis is based on the presence of two or more criteria that encompass the main clinical manifestations of the disease and that were established by the National Institutes of Health (NIH) Consensus Development Conference [4]. NF-1 is constantly associated with vasculopathy and cerebrovascular abnormalities of pathophysiology that is still not understood [5-7]. Several cases have been reported in children, but the incidence of cerebrovascular diseases (CVD) associated with complications and the long-term impacts on these individuals is still poorly recognized, which warrants more studies on the topic [8].

There are multiple case reports in the literature of individuals with NF-1 that developed cerebrovascular diseases (CVD) [9] and heterogeneous neurological manifestations and several complications are present in this association [10], which generates great morbidity and mortality in patients with this genetic condition [11]. The association of NF-1 with CVD is described to result in a range of pathologies, such as cerebral ischemia, aneurysms and *Moyamoya* Syndrome (MMS). The latter, in turn, is characterized by progressive stenosis or occlusion of the internal carotid artery and its branches. For this reason, although lack of full knowledge regarding the natural history, symptoms, etiology and management of such syndrome, it is already known that routine vascular screening/assessment in NF-1 patients is necessary for early identification of this condition [12]. Using such approach, CVD in patients with NF-1 can be diagnosed since childhood. However, this diagnosis is often delayed, as not all children with NF-1 undergo neuroimaging tests [13]. Furthermore, CVDs are still one of the major causes of death, with almost six million people annually [14], aside from being associated with significant morbidity worldwide, which makes them an important public health problem [15]. When CVD are detected early, proper clinical management is able to minimize its impact on adult life [11].

Therefore, detailed studies on the association between NF-1 and CVD are still needed [16]. In this present study, we performed a systematic review to summarize the body of evidence on the association between NF-1 and the development of CVD in children. Increasing knowledge in this field can drive development of more effective protocols to optimize diagnosis and therapy to reduce both mortality and the number of hospitalizations of these patients.

## MATERIALS AND METHODS

### Ethics statement

There were no patients directly involved in the research. The present study used publicly data from previously published studies to perform a systematic review. All information given to the research team was de-identified. Thus, the study was exempted from revision by the Institutional Review Board of the Instituto Gonçalo Moniz, Fundação Oswaldo Cruz, Salvador, Brazil, and did not require signed consent forms.

### Search Strategy

A systematic review of NF-1 and its association with CVD in children aged between 0 to 18 years old was performed, in accordance with the recommendations of the Preferred Reporting Items for Systematic Reviews and Meta-Analyses (PRISMA) report. Two independent reviewers (BBD and FHAG) conducted the research in the following databases: PubMed and EMBASE

The keywords used in the research were: “neurofibromatosis type 1”, “recklinghausen disease”, “children”, “adolescents”, “stroke”, “moyamoya disease”, “vascular diseases”, “cerebrovascular disorders”, “aneurysm”, “congenital abnormalities”. The search strategy used in PubMed and EMBASE is presented in the supplementary file 1. All articles published in English were searched, without time or population restriction of the study. Initially, titles and abstracts were read by the reviewers (Fig 1) and systematic reviews, letters to the editor, case reports and comments were excluded. We did an additional manual search of the reference lists from selected articles to identify eligible publications that may not have appeared in our electronic search strategy. The search was conducted on May 15, 2020. This review was registered in the PROSPERO International Registry (registration number: CRD42020180942).

**Fig 1:**
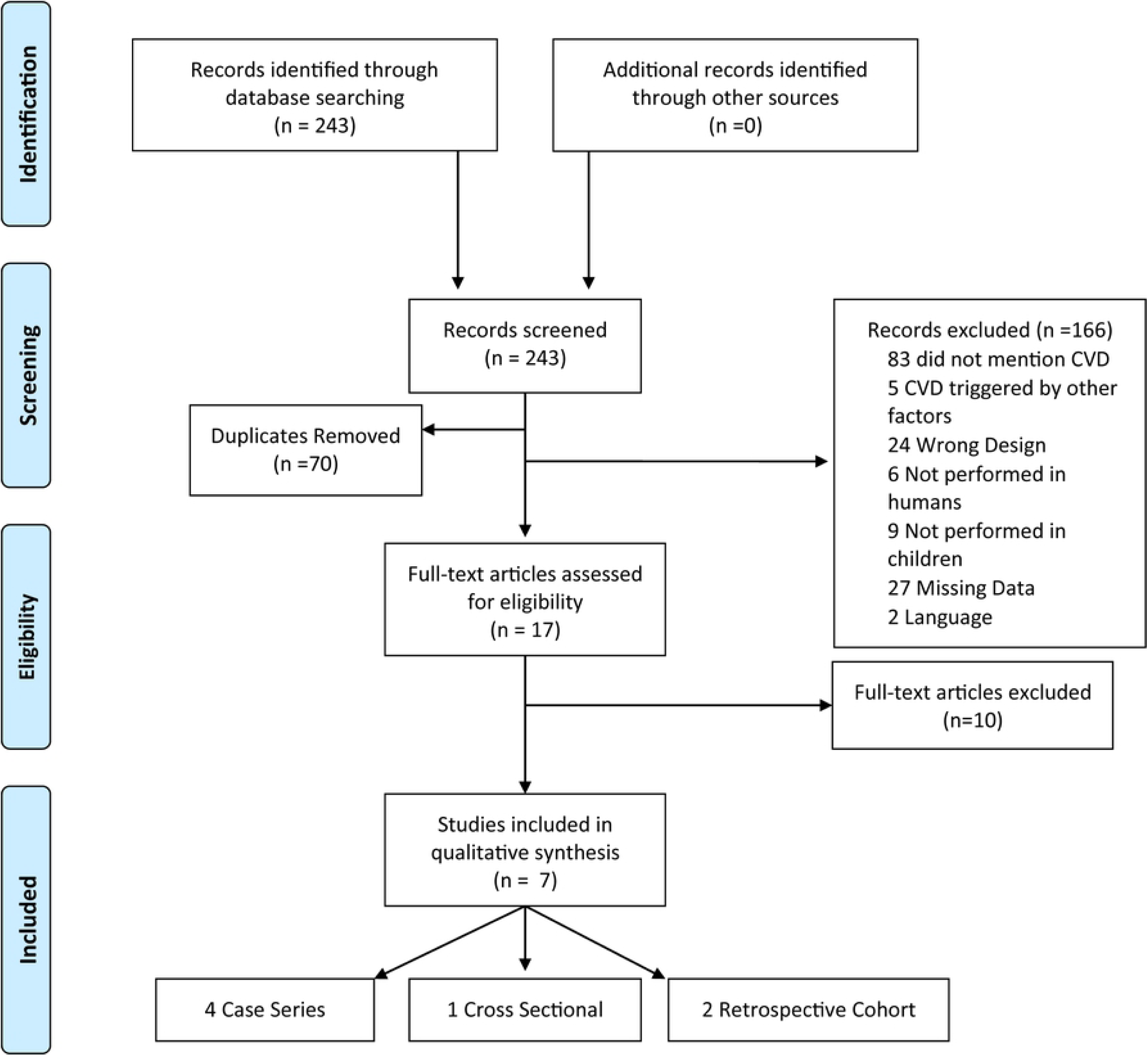
Flowchart of the study selection process study characteristics. Abbreviations: CVD: cerebrovascular disease.

### Data Extraction

Inclusion criteria were: (1) cross-sectional, cohort or case-control studies; (2) studies focusing on NF-1 cerebrovascular disorders; (3) studies carried out with children (from 0 to 18 years old) and (4) studies in which the diagnostic criteria for NF-1 recommended by the NIH were used. Articles in a language other than English, duplicated or not yet published, as well as studies that were not about CVD in patients with NF-1 or that presented cerebrovascular events associated with trauma, neoplasia, radiation treatment or use of medications were excluded. The selection of the studies was divided into 4 steps: 1 reading of the titles, (2) reading of the article abstracts, (3) evaluation of the full articles selected from the previous step and inclusion of other studies present in the reference lists of the selected articles, (4) selection of the studies to include in the systematic review. Data extraction was performed independently by two authors (BBD and FHAG) and the discrepancies between the reviewers were resolved by consensus after discussion with more experienced authors (MBAG, BBA). A table for data extraction was built by each reviewer before writing the manuscript. The table included information on all relevant variables in each retrieved study.

### Data Analysis

All studies addressed the following variables to assess the relationship between NF-1 and CVD in children: age at diagnosis of NF-1, age at the time of image examination to identify CVD, cerebrovascular alteration found, affected vessels, method used for the diagnosis of NF-1, type of image examination performed, associated clinical manifestations and neurosurgery indication(Table 1).

**Table 1.** Summary and characteristics of the articles.

| Study (author, year) | Country | Type of study | Sample size | Age of patients at NF1 diagnosis | Age of patients at image study | Cerebrovascular alteration | Affected vessel | Image study used | Diagnosis os NF1 | Clinical manifestations | Neurosurgery |
| --- | --- | --- | --- | --- | --- | --- | --- | --- | --- | --- | --- |
| (Rosser et al., 2004) | United States of America | Case Series | Total (353);<br>MRI (316);<br>Vasculopathy (8) | Mean, 1.4 years | Mean, 7.3 years | Stenosis / occlusion, cerebral ischemia, ectasia, Moyamoya, aneurysm and hypoplasia | PCA, ICA and MCA | MRI, MRA and Conventional Angiography | NIH diagnostic criteria<sup>a</sup> | Hemiparesis (1), seizure (1) and asymptomatic (7) | 3 patients (exploratory surgery, encephaloduroarteriosynangioses and pial synangiosis) |
| (Cairns et al, 2008) | Australia | Case series | Total (698);<br>MRI (144) ;<br>Vasculopathy (7) | Ranged from 18 months to 5 years | Mean, 6.8 years | Hypoplasia, stenosis/occlusion, collateral vessels | ACA, MCA, PCoA, ICA and PCA | MRI, MRA and Conventional Angiography | NIH diagnostic criteria<sup>a</sup> | Asymptomatic (5), seizure (1), hemiparesis (1), paraesthesia (1) and TIA (1) | 1 patient (pial synangiosis) |
| (Rea et al, 2009) | Canada | Retrospective Cohort | Total (419) ;<br>MRI (266);<br>Arteriopathy (17) | Median, 2 years | Median, 5.2 years (7 children)<br>Median, 5.3 years (10 children) | Stenosis / occlusion, aneurysm, Moyamoya | ICA, MCA and ACA | MRI, MRA, CT, angio-CT and Conventional Angiography | NIH diagnostic criteria<sup>a</sup> | Seizure (3), hemiparesis (1), speech delay (2), Bell's palsy (1), learning disability (4), paraesthesia (1), weakness (2), hyperreflexia (2), ADD (1), hyperphagia (1), episodic dysphasia (1), infantile spasms (1), hypertonia (1), clonus (1), ptosis (1), RAPD (1), hemibalism (1), dystonia (1), aphasia (1), AIT (1) and fine motor delay (1). | 6 patients (pial synangiosis) |
| (Ghosh et al, 2012) | United States of America | Case Series | Total (398);<br>MRI (312);<br>MRA (143);<br>Vasculopathy (15) | Mean, 4.3 ± 3.5 years | Mean, 11.7 SD ± 7.3 years | Stenosis/occlusion, cerebral ischemia, Moyamoya | ICA, MCA, PCA and VA | MRI, MRA, angio-CT and Conventional Angiography | NIH diagnostic criteria<sup>a</sup> | Headache (5), seizure (2) and asymptomatic (7) | 1 patient (encephaloduroarteriomyosynangiosis procedure) |
| (Kaas et al, 2012) | United States of America | Cross Sectional | Total (181);<br>Neuroimaging (77); cerebral vasculopathy (12) | Median 8 years (n=14) | Ranged from 3 to 13 years | Moyamoya, stenosis/occlusion, tortuosity, elongation, displacement, developmental venous anomaly | ACA, MCA, ICA, PCA and VA | MRI, MRA, Conventional Angiography | NIH diagnostic criteria<sup>a</sup> | Weakness (1), reduced responsiveness (1) and asymptomatic (10) | 3 patients (2 - revascularization surgery) |
| (Han et al, 2014) | China | Case Series | Total (6) | Median, 2.7 ± 2.1 years | Mean, 11.4 SD ± 8.3 years | Moyamoya, stenosis / occlusion, intracerebral hemorrhage | ICA, MCA and ACA | MRI, MRA, DSA, SPECT (Conventional digital subtraction angiography, Brain perfusion single-photon emission computerised tomography) | NIH diagnostic criteria<sup>a</sup> | TIA (3), headache (1), intracerebral hemorrhage (1) and cerebral ischemia (1) | 5 patients (Revascularization surgery) |
| (Santoro et al, 2017) | Italy/France | Retrospective cohort | Total (18) | Mean, 2.93 SD± 3.03 years | Mean, 7.43 SD ± 4.27 years | Stenosis/occlusion | MCA, ACA and PCA | MRI, MRA and DSA | NIH diagnostic criteria<sup>a</sup> | Hemiparesis (2), seizure (2), headache (6) and asymptomatic (8) | 11 patients (revascularization surgery) |

**Table Note:** <sup>a</sup>National Institutes of Health (NIH) Consensus Development Conference diagnostic criteria: consists of the presence of two or more of the following characteristics - six or more cafe au lait macules over 5 mm in greatest diameter in prepubertal individuals and over 15 mm in greatest diameter in post pubertal individuals; two or more neurofibromas of any type or one plexiform neurofibroma; freckling in the axillary or inguinal regions; optic glioma; two or more Lisch nodules (iris
hamartomas); a distinctive osseous lesion such as sphenoid dysplasia or thinning of long bone cortex, with or without pseudarthrosis and a first-degree relative (parent, sibling, or offspring) with NF-1 by the above criteria.
**Abbreviations:** NF-1:Neurofibromatosis Type 1; PCA: posterior cerebral artery; MCA: middle cerebral artery; ACA: anterior cerebral artery; ICA: internal carotid artery; PCoA: posterior communicating artery; VA: vertebral artery; MRI: magnetic resonance imaging; MRA: magnetic resonance angiography; CA: conventional angiography; CT: computed tomography; angio-CT: computed tomography angiography; DSA: digital subtraction angiography; SPECT: Brain perfusion single-photon emission computerized tomography; TIA: transient ischemic attack; RAPD: DPAR: relative afferent pupillary defect; ADD: attention deficit disorder; SD: standard variation

### Quality Assessment

The methodological quality of the studies included in the meta-analysis was assessed using the Newcastle-Ottawa Scale (NOS) (Table 2) [17]. NOS scores of 0–3, 4–6 and 7–9 were considered to be low, moderate and high quality, respectively. The overall methodological quality was then summarized based on the observed quality trends.

**Table 2:**
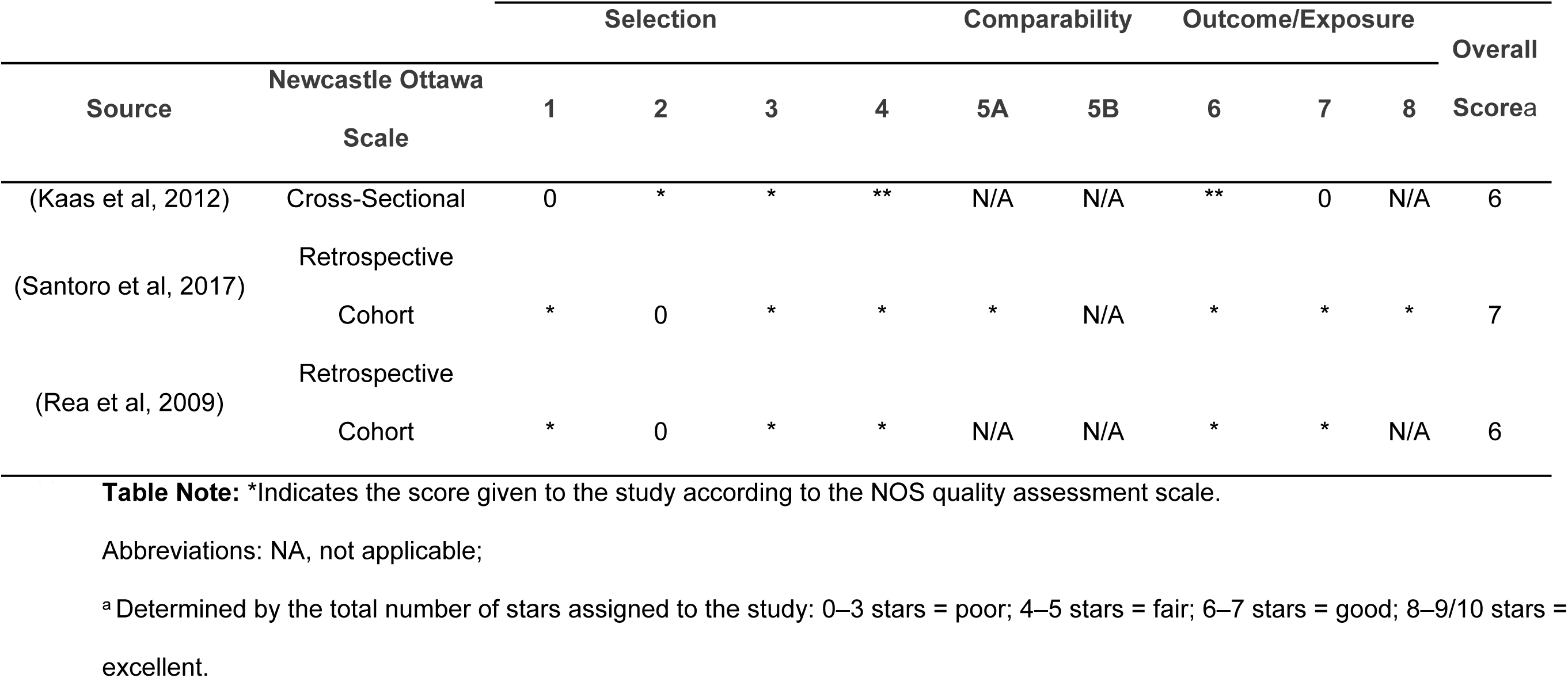
Quality Assessment of Studies Included in the Systematic Review by Newcastle Ottawa Scale.

## RESULTS

### Study characteristics and quality evaluation

Initially, 243 studies were selected from the main database search. After detailed review, 70 duplicates were removed and 166 articles were excluded: 83 did not address CVD, 5 addressed CVD triggered by other factors, such as a particular medicine or neoplasms, 18 were case reports, 4 were letters to the editor and 2 were brief communications (Figure 1). Subsequently, 6 other studies were excluded for not being performed in humans, 9 for not being performed in children, 27 were not found or published and 2 were in a language other than English (Polish and German) (Fig 1). Finally, 10 studies that did not provide sufficient clinical data on the pediatric population were still excluded. The remaining studies (7), which describe NF-1 associated with CVD in children, were included in this review. Fig 1 summarizes the article selection process.

Among the seven selected studies, 2 had a medical record review project (retrospective cohort) 1 was a cross-sectional study and 4 were series of cases. In all of them, the NF-1 diagnostic criterion used was that developed by the NIH. Moreover, the eligible studies (cohort and cross-sectional) for the evaluation of the Newcastle-Ottawa Scale (NOS) were of good quality presenting between 6-7 stars (Table 2). Such studies were performed out in six different countries: one in Australia, one in Canada, one in China, one in Italy and France and three of them in different cities in the United States of America (Washington DC, Cleveland and Baltimore) (Fig 2).

**Fig. 2:**
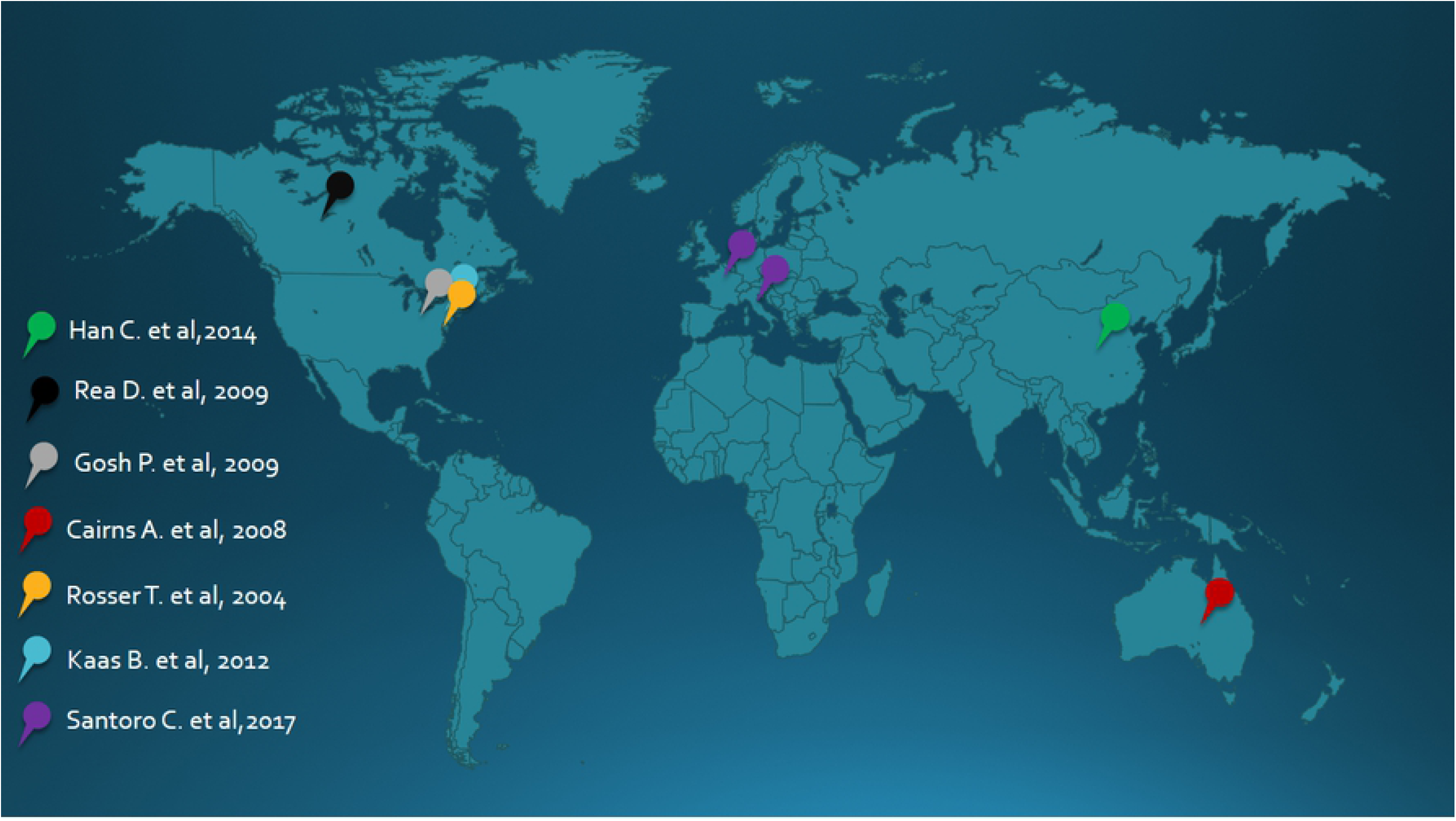
Distribution of studies used around the world.

### Cerebrovascular alterations presented higher frequency at Willis polygon

In this review, we detected 12 types of cerebrovascular changes significantly associated with NF-1 and / or disease progression in children (Table 1). In six of the seven studies the internal carotid artery was the most affected vessel among patients: 75% in the study by Rosser et al., 80% in the study by Ghosh et al., and 100% in the studies by Read et al., Kaas et al., Han et al. and Santoro et al. In the study by Cairns et al. the most affected vessels were those of the Willis polygon (85,7% of patients) (Table 1 and Fig 3).

**Fig. 3:**
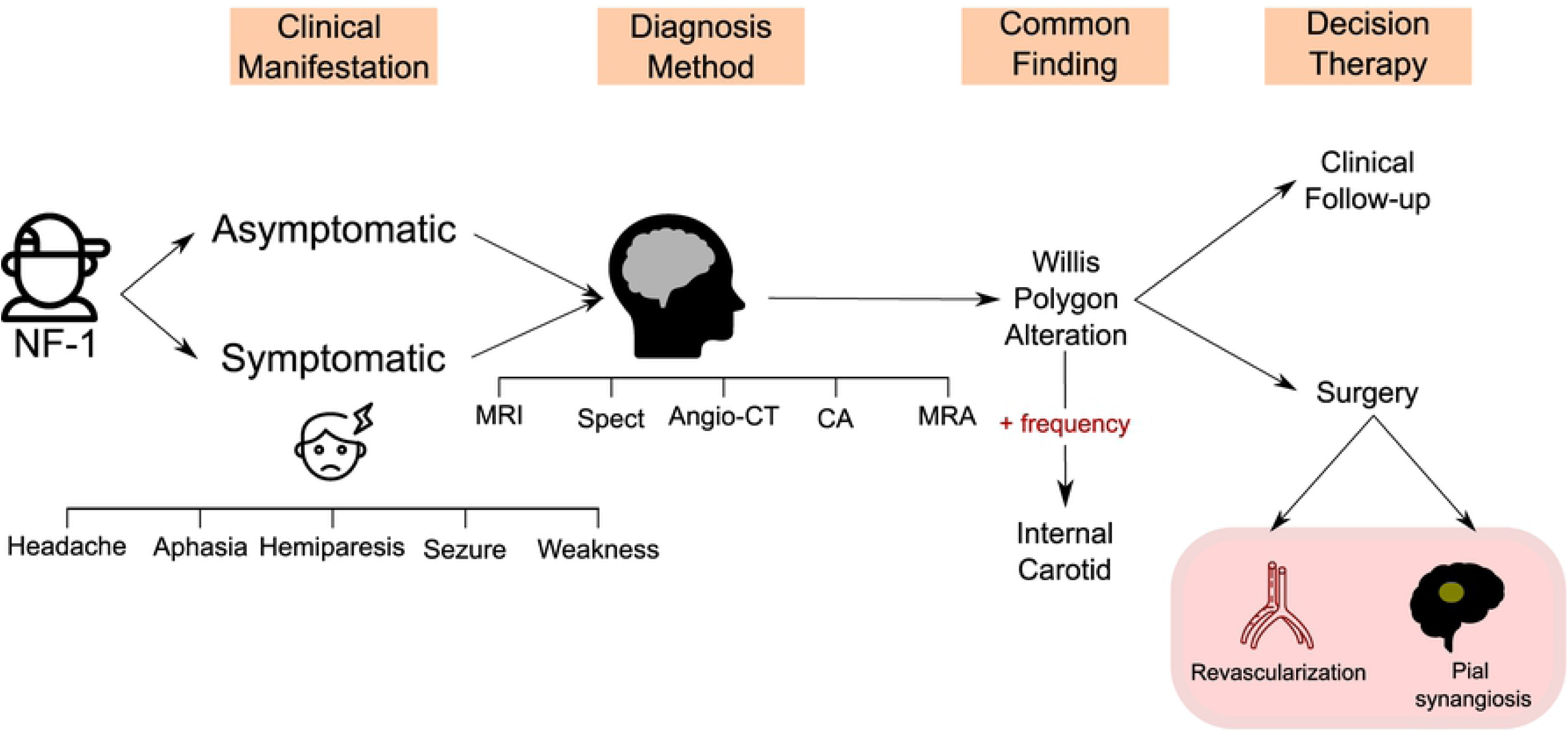
Flow diagram summarizing the main results of the systematic review. Abbreviations: NF-1:Neurofibromatosis Type 1; MRI: magnetic resonance imaging; MRA: magnetic resonance angiography; CA: conventional angiography; CT: computed tomography; angio-CT: computed tomography angiography; SPECT: Brain perfusion single-photon emission computerized tomography.

### Radiologic investigation

The diagnosis of NF-1 was performed prior to brain imaging exams in all studies, showing that many CVDs could have been previously diagnosed, which possibly would prevent complications or more invasive treatments. The most frequently performed imaging tests were Magnetic Resonance Angiography (MRA) and Magnetic Resonance Imaging (MRI) performed in 100% of the studies evaluated followed by Conventional Angiography (CA) - 71.4%, Computed Tomography Angiography (Angio-CT) and Digital Substraction Angiography DSA - 28.5% each and finally Brain Perfusion Single-photon Emission Computerized Tomography (SPECT) and Computed Tomography (CT) - 14.2% each. (Table 1 and Fig 3).

### The heterogeneous spectrum of clinical manifestations and delays in diagnosis of CVD

NF-1 has been closely associated with *MMS* vasculopathy in five studies and its presentation is related to worse outcomes and more evident clinical manifestations. In the study by Rea et al., It was possible to correlate NF-1 with the difficulty of developing cognitive functions such as reading, speaking and the ability to maintain attention. Furthermore, in the study by Ghosh et al, five of the fifteen patients evaluated presented headache as the only clinical manifestation that could indicate CVD. Thus, headache is possibly an important warning sign in this population.

In general, the clinical manifestations found were very heterogeneous (Fig 3). Patients with a history of paresthesia, seizures, and even severe strokes were observed, among other manifestations. However, it is important to note that until the time of diagnosis of CVD, 44,5% of the patients were asymptomatic, which shows the importance of screening in childhood. Furthermore, the type of surgery most reported in the studies evaluated was *pial synangiosis* surgery (five of seven studies) followed by revascularization surgery (three of seven studies) (Fig 3). In the study by Santoro *et al*. patients who underwent surgical treatment (50%) exhibited global clinical stability.

## DISCUSSION

NF-1 is a rare, autosomal dominant disease [18], which has a strong relationship with vascular malformations [19] and whose first symptoms commonly appear in childhood [20]. Such vascular alterations, although they can occur in any part of the child body [8], have a greater severity when presented in the central nervous system (CNS) [10]. For this reason, knowledge of the association between NF-1 and cerebrovascular diseases triggered by this disease can contribute to a better understanding of the pathophysiology of NF-1 manifestations in the CNS, as well as guide development of novel approaches to prevent cerebrovascular events in children with NF-1. This systematic review demonstrated that the use of early imaging methods is associated with early interventions and, consequently, more favorable outcomes, even in previously asymptomatic children.

Our search identified that all studies selected for systematic analysis reported that some artery of the Willis polygon was affected [9, 11-13, 21, 22], of which the internal carotid artery is shown as the main site of alteration among all evaluated patients. Importantly, the branches generated by the internal carotid artery are the middle cerebral artery and anterior cerebral artery [23], responsible for predominantly motor and somatosensory deficits [24]. Thus, the manifestations presented can vary from a focal motor deficit to an aphasia [13], which makes early diagnosis difficult [25] since depending on the age of the child communication is still not effective and motor skills has not well-developed. Therefore, it is crucial that more studies are developed to assess the need for periodic screening in this group through brain imaging.

Of note, approximately 44.5% of the patients were asymptomatic at the time of the imaging exam, which showed vascular changes and rises the need of early screening. At first glance, this fact counts against the NIH NF-1 management manuals since brain imaging is only indicated in the presence of symptoms. In addition, in the study by Ghosh et al, 33.4% of the patients evaluated presented headache as the only symptom, which can be justified by the fact that the most common vascular alteration reported in all studies was arterial stenosis / occlusion. Moreover, the comparison between the selected studies about children with NF-1 and cerebrovascular diseases revealed that the earlier the imaging tests are performed, even when the children are asymptomatic at the time of the examination could contribute to better interventions. This corroborates the hypothesis that the performance of periodic and early examinations can impact on morbidity and mortality and on the quality of life of children with NF-1.

The available evidence allowed us to conclude that neurological manifestations of NF-1 in the pediatric group are heterogeneous and difficult to diagnose clinically, and in most cases, it is necessary to perform a complementary imaging test. In addition, when these manifestations are associated with MMS the children have a worse prognosis [22]. Notably, *MMS* is described with the severity of symptoms presented by children [12], what can be seen in six of the seven studies evaluated. Moreover, the neurosurgery most performed by the studies was pial synangiosis surgery, widely used for the treatment of *MMS* and reduction of its clinical manifestations. An important observation is that the surgical interventions performed were also more successful when associated with early imaging tests, which corroborates the idea of early screening for recognition of vascular malformations on the CNS. The tests most used by the studies evaluated were MRA and MRI, although there is no consensus on which is best test to be performed or the periodicity established for that execution.

Our study has important limitations. Some studies did not contain sufficient information about the pediatric group, not allowing an adequate assessment. In addition, most studies are only descriptive, not performing statistical analysis of the data presented, which made it impossible to perform a meta-analysis. It is also important to highlight the low availability and quality of the studies carried out about this theme, which present a very variable number of patients and a little detailed analysis of each individual patient, which made comparisons between groups difficult. Besides, due to the small amount of evidence available, it was necessary to use the series of cases, which brings an important limitation since we were unable to apply an adequate quality scale to the studies. In addition, the available data on the chronology of the imaging exams and the reason for the indication of one method within the other is very scarce, which makes it difficult to interpret these parameters. Despite this, as it is a rare disease that affects 1 in every 3000 live births [26]and has major complications[10], we believe that this systematic review reaches the proposed ideas.

The results of this systematic review show that the possible implementation of screening measures using imaging methods has the potential to improve the early diagnosis of cerebrovascular alterations in children with NF-1 as well as early intervention. This could potentially reduce the number of deaths and sequelae caused by the diagnosis of vascular change only when an injury is already installed. Even so, more robust and better-quality studies are needed to clarify the frequency and which exams should be recommended within each age group. An evaluation of imaging methods in isolation and in conjunction with clinical parameters is necessary in order to draw a better line of care for these children since birth.

## Acknowledgments

The authors thank to Dr. Carla Daltro or helping with the initial version of this paper.

## Privacy statement

the authors guarantee that all data were anonymized and complies with the Helsinki declaration.

## Availability of data and materials

The datasets used and/or analyzed during the current study are all publicly available.

## REFERENCES

1. Souza JF, Toledo LL, Ferreira MC, Rodrigues LO, Rezende NA. [Neurofibromatosis type 1: more frequent and severe then usually thought]. Rev Assoc Med Bras (1992). 2009;55(4):394-9. Epub 2009/09/15. doi: 10.1590/s0104-42302009000400012. PubMed PMID: 19750304.

2. Evans DG, Howard E, Giblin C, Clancy T, Spencer H, Huson SM, et al. Birth incidence and prevalence of tumor-prone syndromes: estimates from a UK family genetic register service. Am J Med Genet A. 2010;152A(2):327–32. Epub 2010/01/19. doi: 10.1002/ajmg.a.33139. PubMed PMID: 20082463.

3. Friedman JM. Neurofibromatosis 1. In: Adam MP, Ardinger HH, Pagon RA, Wallace SE, Bean LJH, Stephens K, et al., editors. GeneReviews((R)). Seattle (WA)1993.

4. Ferner RE, Gutmann DH. Neurofibromatosis type 1 (NF1): diagnosis and management. Handb Clin Neurol. 2013;115:939-55. Epub 2013/08/13. doi: 10.1016/B978-0-444-52902-2.00053-9. PubMed PMID: 23931823.

5. Gkampeta A, Hatzipantelis E, Kouskouras K, Pavlidou E, Pavlou E. Ischemic cerebral infarction in a 5-year-old male child with neurofibromatosis type 1. Childs Nerv Syst. 2012;28(11):1989-91. Epub 2012/05/10>. doi: 10.1007/s00381-012-1790-0. PubMed PMID: 22570170.

6. Yilmaz H, Erkin G, Gumus H, Nalbant L. Coexistence of Neurofibromatosis Type-1 and MTHFR C677T Gene Mutation in a Young Stroke Patient: A Case Report. Case Rep Neurol Med. 2013;2013:735419. Epub 2013/03/28. doi: 10.1155/2013/735419. PubMed PMID: 23533858; PubMed Central PMCID: PMCPMC3600223.

7. Terry AR, Jordan JT, Schwamm L, Plotkin SR. Increased Risk of Cerebrovascular Disease Among Patients With Neurofibromatosis Type 1: Population-Based Approach. Stroke. 2016;47(1):60-5. Epub 2015/12/10. doi: 10.1161/STROKEAHA.115.011406. PubMed PMID: 26645253.

8. Sobata E, Ohkuma H, Suzuki S. Cerebrovascular disorders associated with von Recklinghausen’s neurofibromatosis: a case report. Neurosurgery. 1988;22(3):544-9. Epub 1988/03/01. doi: 10.1227/00006123-198803000-00016. PubMed PMID: 3129670.

9. Cairns AG, North KN. Cerebrovascular dysplasia in neurofibromatosis type 1. J Neurol Neurosurg Psychiatry. 2008;79(10):1165-70. Epub 2008/05/13. doi: 10.1136/jnnp.2007.136457. PubMed PMID: 18469031.

10. Giannantoni NM, Broccolini A, Frisullo G, Pilato F, Profice P, Morosetti R, et al. Neurofibromatosis type 1 associated with vertebrobasilar dolichoectasia and pontine ischemic stroke. J Neuroimaging. 2015;25(3):505-6. Epub 2014/09/19. doi: 10.1111/jon.12160. PubMed PMID: 25230986.

11. Ghosh PS, Rothner AD, Emch TM, Friedman NR, Moodley M. Cerebral vasculopathy in children with neurofibromatosis type 1. J Child Neurol. 2013;28(1):95-101. Epub 2012/04/26. doi: 10.1177/0883073812441059. PubMed PMID: 22532547.

12. Santoro C, Di Rocco F, Kossorotoff M, Zerah M, Boddaert N, Calmon R, et al. Moyamoya syndrome in children with neurofibromatosis type 1: Italian-French experience. Am J Med Genet A. 2017;173(6):1521-30. Epub 2017/04/20. doi: 10.1002/ajmg.a.38212. PubMed PMID: 28422438.

13. Rea D, Brandsema JF, Armstrong D, Parkin PC, deVeber G, MacGregor D, et al. Cerebral arteriopathy in children with neurofibromatosis type 1. Pediatrics. 2009;124(3):e476-83. Epub 2009/08/27. doi: 10.1542/peds.2009-0152. PubMed PMID: 19706560.

14. Organization WH. The global burden of cerebrovascular disease 2000.

15. Dobkin B. The economic impact of stroke. Neurology. 1995;45(2 Suppl 1):S6-9. Epub 1995/02/01. PubMed PMID: 7885589.

16. Zhang H, Zheng L, Feng L. Epidemiology, diagnosis and treatment of moyamoya disease. Exp Ther Med. 2019;17(3):1977-84. Epub 2019/03/15. doi: 10.3892/etm.2019.7198. PubMed PMID: 30867689; PubMed Central PMCID: PMCPMC6395994.

17. Stang A. Critical evaluation of the Newcastle-Ottawa scale for the assessment of the quality of nonrandomized studies in meta-analyses. Eur J Epidemiol. 2010;25(9):603-5. Epub 2010/07/24. doi: 10.1007/s10654-010-9491-z. PubMed PMID: 20652370.

18. Mahalingam M. NF1 and Neurofibromin: Emerging Players in the Genetic Landscape of Desmoplastic Melanoma. Adv Anat Pathol. 2017;24(1):1–14. Epub 2016/12/13. doi: 10.1097/PAP.0000000000000131. PubMed PMID: 27941538.

19. Rosser TL, Vezina G, Packer RJ. Cerebrovascular abnormalities in a population of children with neurofibromatosis type 1. Neurology. 2005;64(3):553–5. Epub 2005/02/09. doi: 10.1212/01.WNL.0000150544.00016.69. PubMed PMID: 15699396.

20. Chisholm AK, Anderson VA, Pride NA, Malarbi S, North KN, Payne JM. Social Function and Autism Spectrum Disorder in Children and Adults with Neurofibromatosis Type 1: a Systematic Review and Meta-Analysis. Neuropsychol Rev. 2018;28(3):317–40. Epub 2018/08/12. doi: 10.1007/s11065-018-9380-x. PubMed PMID: 30097761.

21. Kaas B, Huisman TA, Tekes A, Bergner A, Blakeley JO, Jordan LC. Spectrum and prevalence of vasculopathy in pediatric neurofibromatosis type 1. J Child Neurol. 2013;28(5):561–9. Epub 2012/07/27. doi: 10.1177/0883073812448531. PubMed PMID: 22832780; PubMed Central PMCID: PMCPMC3496801.

22. Han C, Yang WZ, Zhang HT, Ye T, Duan L. Clinical characteristics and long-term outcomes of moyamoya syndrome associated with neurofibromatosis type 1. J Clin Neurosci. 2015;22(2):286–90. Epub 2014/12/03. doi: 10.1016/j.jocn.2014.05.046. PubMed PMID: 25443089.

23. Almeida JP, Reghin Neto M, Chaddad Neto F, E DEO. Anatomical considerations in the treatment of intracranial aneurysms. J Neurosurg Sci. 2016;60(1):27–43. Epub 2015/12/15. PubMed PMID: 26657135.

24. Defebvre L, Krystkowiak P. Movement disorders and stroke. Rev Neurol (Paris). 2016;172(8-9):483–7. Epub 2016/08/02. doi: 10.1016/j.neurol.2016.07.006. PubMed PMID: 27476417.

25. Cnossen MH, Moons KG, Garssen MP, Pasmans NM, de Goede-Bolder A, Niermeijer MF, et al. Minor disease features in neurofibromatosis type 1 (NF1) and their possible value in diagnosis of NF1 in children < or = 6 years and clinically suspected of having NF1. Neurofibromatosis team of Sophia Children’s Hospital. J Med Genet. 1998;35(8):624–7. Epub 1998/08/27. doi: 10.1136/jmg.35.8.624. PubMed PMID: 9719365; PubMed Central PMCID: PMCPMC1051384.

26. Alves Junior SF, Zanetti G, Alves de Melo AS, Souza AS, Jr., Souza LS, de Souza Portes Meirelles G, et al. Neurofibromatosis type 1: State-of-the-art review with emphasis on pulmonary involvement. Respir Med. 2019;149:9–15. Epub 2019/03/20. doi: 10.1016/j.rmed.2019.01.002. PubMed PMID: 30885426.

